# Dietary Lees Supplementation Enhances Poultry Health Performance Through Gut Ecosystem Reprogramming

**DOI:** 10.64898/2026.01.23.700869

**Authors:** M. Shaminur Rahman, Suborna Islam, Tamanna Jerin Anannya, Adnan Muyeed, Moumita Rahman Sazza, Mohammad Imtiaj Uddin Bhuiyan, Selina Akter

## Abstract

The search for sustainable alternatives to in-feed antibiotics has intensified with the global antimicrobial resistance crisis. While microbial by-products show potential, their mechanisms remain elusive. This study reveals that spent yeast lees, a fermentation by-product, profoundly reprograms the chicken gut ecosystem to enhance growth. We investigated the differential effects of live spent lees versus autoclaved spent lees from a molasses fermenter with *Saccharomyces cerevisiae*. In a 14-day controlled trial, 0-day-old poultry chicks (n=15) were assigned to a control diet (n=5), a live spent Lees-supplemented diet (n=5), or an autoclaved spent Lees-supplemented diet (n=5). Through the feeding trial with either live spent Lees or autoclaved spent Lees, we demonstrate significant enhancement of growth performance, with autoclaved spent Lees achieving the highest final body weight (573.4 ± 17.2 g). Using deep shotgun metagenomic sequencing of faecal samples, we demonstrate that both spent Lees forms induced a dramatic ecological shift, driving the gut microbiome to a state of monodominance by *Bacteroides fragilis* (reaching 74.83% with live spent Lees). This was accompanied by the near-complete eradication of enteropathogens like like *Enterococcus faecium* and *Escherichia coli*. Crucially, live spent Lees supplementation effectively countered a natural age-dependent expansion of the gut resistome, eliminating the high-risk oxazolidinone resistance gene *o23S* and restructuring the virulome away from offensive effector systems. However, this beneficial restructuring came at the cost of significantly reduced microbial diversity and suppression of beneficial commensals. A key finding was that autoclaved spent Lees, while inducing a less extreme microbial shift, yielded superior final body weight, suggesting its lysed cells provide enhanced nutritional availability. Our work establishes spent yeast Lees not as mere waste, but as a potent, multi-functional supplement that enhances growth through gut ecosystem engineering, yet necessitates a strategic choice between its live and autoclaved forms based on production priorities.

## 1. Introduction

Antimicrobial resistance (AMR) significantly exacerbated by the extensive use of in-feed antibiotics in livestock, necessitates an urgent focus on sustainable, non-antibiotic feed alternatives ^1,2^. The poultry industry is under immense pressure to identify supplements that can effectively enhance animal health performance (growth, welfare, disease resistance) while mitigating the development and transfer of antibiotic resistance genes within the gut resistome^3^.

Spent yeast lees, the residual biomass from industrial fermentation processes (*Saccharomyces cerevisiae* from molasses fermenters), represent a high-volume, cost-effective by-product poised for valorization. These lees are rich in proteins, nucleotides, and bioactive cell wall components like β-glucans and mannan-oligosaccharides^4^. Research on similar fermentation by-products, such as wine lees and sake lees, demonstrates their multifunctional potential beyond basic nutrition. Lees and vinification by-products have been shown to improve antioxidant status and modulate immune-related gene expression in broilers, suggesting they can boost the animal’s defense mechanisms and protect product quality^5^. For instance, wine lees can mitigate oxidative stress, enhance egg production and yolk lipid stability with extending its shelf life^6,7^. A different research found Sake lees to specifically improve intestinal barrier function in meat-type chickens by increasing tight junction protein expression^8^. While these studies confirm the broad health and nutritional potential of yeast by-products, a crucial mechanistic gap remains. There is a lack of high-resolution data demonstrating how complex supplements like spent lees, especially when differentially processed (live versus autoclaved), exert their effects on the genomic profile and ecological structure of the avian gut microbiome. Critically, we do not fully understand the precise mechanism by which these supplements influence mobile AMR genes that are commonly found in poultry-associated bacteria^1,2^. We conducted a controlled trial experiments on broiler chicks and employed deep shotgun metagenomic sequencing. This work establishes spent yeast lees as a potent, multifunctional supplement that achieves growth enhancement through gut ecosystem reprogramming.

## 2. Methodology

### 2.1 Ethical Approval

The study protocol was approved by the Ethical Review Committee of the Faculty of Biological Sciences, Jashore University of Science and Technology, Jashore, Bangladesh (Approval No.: ERC/FBST/JUST/2025-261). All experimental procedures were conducted in accordance with international ethical standards, including the ARRIVE guidelines (Animal Research: Reporting of In Vivo Experiments)^9^. Continuous veterinary supervision was ensured to maintain adherence to these ethical standards throughout the study.

### 2.2 Study Design and Feeding Regimen

An experimental study was conducted over a 14-day period to investigate the effects of dietary yeast supplementation on SPF (specific pathogen free) poultry, specifically Hubbard Classic broiler chickens from Afil Poultry Farms. A total of fifteen (n=15) 0 day-old poultry chicks (*Gallus gallus domesticus*) were randomly allocated into three dietary treatment groups: (1) a Control group (n=5) fed a basal diet only; (2) a Spent Lees group (n=5) fed the basal diet supplemented with live yeast culture (n=5); and (3) an Autoclaved Spent Lees group (n=5) fed the basal diet supplemented with autoclaved (heat-inactivated) yeast. The basal feed composition is detailed in Supplementary Table S1. The yeast spent Lees, which is a mainly solid yeast-rich residue from the bottom of the fermenter, was sourced from the Carew & Co (Bangladesh) Ltd. It is a distillery located in Darshana, uses large tanks to ferment molasses for the production of spirits. The concentration of live *Saccharomyces cerevisiae* (Yeast) cells in the spent Lees was approximately 4 × 10 CFU/mL. The feeding regimen was progressively increased throughout the study to meet the birds’ growth requirements. The control group received a daily ration of basal feed, starting from 50g on day 1 and increasing by 25g every day to 375g on day 14. The yeast-supplemented groups received the same daily quantity of basal feed, with an additional liquid yeast supplement. The supplement was incorporated into the basal feed at a rate of 10% (v/w), beginning with 5 mL of spent Lees on day 1 and gradually increasing to 37.5 mL by day 14 (Table 1). Individual body weights were recorded for all chickens at the start of the experiment (Day 0), on Day 7, and at the conclusion of the study (Day 14) (Table 2).

**Table 1:** Daily Feeding Regimen Over the 14-Day Experimental Period.

| Days | Control (Basal feed) | Basal feed + Spent Lees | Basal feed + Autoclaved Spent Lees |
| --- | --- | --- | --- |
| 1 | 50g | 50g + 5ml | 50g + 5ml |
| 2 | 75g | 75g + 7.5ml | 75g + 7.5ml |
| 3 | 100g | 100g + 10ml | 100g + 10ml |
| 4 | 125g | 125g + 12.5ml | 125g + 12.5ml |
| 5 | 150g | 150g + 15ml | 150g + 15ml |
| 6 | 175g | 175g + 17.5ml | 175g + 17.5ml |
| 7 | 200g | 200g + 20ml | 200g + 20ml |
| 8 | 225g | 225g + 22.5ml | 225g + 22.5ml |
| 9 | 250g | 250g + 25ml | 250g + 25ml |
| 10 | 275g | 275g + 27.5ml | 275g + 27.5ml |
| 11 | 300g | 300g + 30ml | 300g + 30ml |
| 12 | 325g | 325g + 32.5ml | 325g + 32.5ml |
| 13 | 350g | 350g + 35ml | 350g + 35ml |
| 14 | 375g | 375g + 37.5ml | 375g + 37.5ml |

**Table 2:** Effects of Spent Lees and Autoclaved Spent Lees on Body Weight in Poultry Chicken.

| Days | Control | Average ± SD | Live Yeast | Average ± SD | Autoclaved Yeast | Average ± SD |
| --- | --- | --- | --- | --- | --- | --- |
| Day 0 | 102 | 98.4 ± 5.0 | 96 | 86.2 ± 11.4 | 83 | 90.2 ± 5.1 |
|  | 98 |  | 68 |  | 95 |  |
|  | 98 |  | 86 |  | 88 |  |
|  | 90 |  | 96 |  | 93 |  |
|  | 104 |  | 85 |  | 92 |  |
| Day 7 | 219 | 213.4 ± 6.1 | 265 | 251.4 ± 21.4 | 225 | 239.6 ± 11.4 |
|  | 209 |  | 283 |  | 253 |  |
|  | 216 |  | 228 |  | 238 |  |
|  | 205 |  | 241 |  | 235 |  |
|  | 218 |  | 240 |  | 247 |  |
| Day 14 | 545 | 526.2 ± 18.6 | 620 | 545.4 ± 66.3 | 550 | 573.4 ± 17.2 |
|  | 507 |  | 450 |  | 597 |  |
|  | 540 |  | 547 |  | 575 |  |
|  | 503 |  | 530 |  | 575 |  |
|  | 536 |  | 580 |  | 570 |  |

### 2.3 DNA Extraction and Sample Pooling

At Day 0 and Day 14, fecal samples were collected from individual chickens for microbial community analysis. Genomic DNA was extracted from approximately 200 mg of each sample using the QIAamp® Fast DNA Stool Mini Kit (Qiagen, catalog no. 51604) according to the manufacturer’s instructions. The extracted DNA was eluted in a final volume of 50 µl of elution buffer. The concentration of each DNA extract was quantified, and samples were subsequently pooled by treatment group and time point to create representative composite samples for downstream sequencing, as detailed in Table 3. The pooling strategy was designed to normalize the contribution of each individual sample based on its DNA concentration. Specifically, the volume of each individual DNA extract contributed to a pool was inversely proportional to its concentration, ensuring an equimolar representation. This resulted in the creation of seven pooled libraries: one for the Control group at Day-0 (CP-1), two for the Control group at Day-14 (CP-2, CP-3), two for the Live Yeast group at Day-14 (LYP-4, LYP-5), and two for the Autoclaved Yeast group at Day-14 (AYP-6, AYP-7). Each final pool had a total volume of 40 µl and a quantified concentration suitable for subsequent shotgun metagenomic sequencing (Table 3).

**Table 3:** DNA Extraction Metrics and Sample Pooling Strategy by Treatment Group.

| Day | Treatment | Sample ID<br>(50µl Elution) | DNA Conc.<br>(ng/µl) | Pool amount<br>(µl) | Pool_ID | Final Conc<br>(ng/µl) | Total Volume<br>(30-40 µl) | DNA Extraction<br>Kit Name with<br>Catalog ID |
| --- | --- | --- | --- | --- | --- | --- | --- | --- |
| Day-0 | Control | 1 | 3.3 | 10.5 | CP-1 | 3.46 | 40 µl | QIAamp® Fast<br>DNA Stool Mini Kit<br>(cat. no. 51604) |
| Day-0 | Control | 2 | 3.94 | 8.8 |  |  |  |  |
| Day-0 | Control | 3 | 4.46 | 7.8 |  |  |  |  |
| Day-0 | Control | 4 | 2.66 | 13.0 |  |  |  |  |
| Day-0 | Control | 5 | 0.78 | 10.5 |  |  |  |  |
| Day-14 | Control | 6 | 19.8 | 5.8 | CP-2 | 8.62 | 40 µl |  |
| Day-14 | Control | 7 | 10.9 | 10.5 |  |  |  |  |
| Day-14 | Control | 8 | 4.86 | 23.7 |  |  |  |  |
| Day-14 | Control | 9 | 27.2 | 13.1 | CP-3 | 17.86 | 40 µl |  |
| Day-14 | Control | 10 | 13.3 | 26.9 |  |  |  |  |
| Day-14 | Spent Lees | 11 | 2.14 | 22.4 | LYP-4 | 3.6 | 40 µl |  |
| Day-14 | Spent Lees | 12 | 2.98 | 16.1 |  |  |  |  |
| Day-14 | Spent Lees | 13 | 32.4 | 1.5 |  |  |  |  |
| Day-14 | Spent Lees | 14 | 3.02 | 22.2 | LYP-5 | 3.35 | 40 µl |  |
| Day-14 | Spent Lees | 15 | 3.76 | 17.8 |  |  |  |  |
| Day-14 | Autoclave Spent Lees | 16 | 3.48 | 30.8 | AYP-6 | 8.05 | 40 µl |  |
| Day-14 | Autoclave Spent Lees | 17 | 43.6 | 2.5 |  |  |  |  |
| Day-14 | Autoclave Spent Lees | 18 | 15.9 | 6.8 |  |  |  |  |
| Day-14 | Autoclave Spent Lees | 19 | 35.4 | 4.3 | AYP-7 | 7.61 | 40 µl |  |
| Day-14 | Autoclave Spent Lees | 20 | 4.26 | 35.7 |  |  |  |  |

### 2.4 Library Preparation and Shotgun Metagenomic Sequencing

Shotgun metagenomic libraries were constructed from the seven pooled DNA samples following the iNextEra library preparation protocol, adapted from Jones et al. 2023^10^ and Bhuiyan et al. 2025^11^. The application of this protocol has been published in our other research paper^12^. The process began with tagmentation of DNA using Illumina bead-linked transposomes (Illumina DNA library prep, Illumina, Inc., San Diego, CA, USA) combined with 2X TMP buffer in a 6 µL reaction containing 3 µL of 2X TMP buffer, 0.5 µL of Bead-linked transposome, and 2.5 µL of DNA template, followed by incubation at 53°C for 30 minutes. A limited-cycle amplification was then conducted to incorporate full-length Illumina adapters with dual indices, employing custom iNextEra primers designed for multiplexed sequencing. The PCR was set up by first preparing a 51.5 µL master mixture containing 40.25 µL HyClone water (Cytiva Life Sciences, Marlborough, MA, USA), 9.5 µL of Q5 Reaction Buffer (New England Biolabs, Ipswich, MA, USA), 1.25 µL of 10mM dNTPs (Roche Diagnostic Corporation, Indianapolis, IN, USA), and 0.5 µL of Q5 High-Fidelity DNA Polymerase (New England Biolabs, Ipswich, MA, USA). This master mixture was transferred to the 6 µL tagmentation product, after which 2.5 µL of 5 µM iNextEra5 index primer and 2.5 µL of 5 µM iNextEra7 index primer were incorporated, bringing the final reaction volume to 62.5 µL with unique barcode combinations. Thermal cycling was executed on a T100 instrument (Bio-Rad Laboratories, Inc., Hercules, CA, USA) using the following parameters: initial incubation at 72°C for 3 minutes for end-repair; polymerase activation at 98°C for 3 minutes; 12 amplification cycles of 98°C for 45 seconds, 62°C for 30 seconds, and 72°C for 2 minutes; with a terminal extension at 72°C for 1 minute. Amplification success and library size distribution were verified by resolving 5 µL of PCR product on an agarose gel, confirming a fragment smear within the 300-700 bp target range. Final library processing involved quantification with the Qubit Flex system (Thermo Fisher Scientific, Waltham, MA USA), followed by normalization, pooling, and purification using AMPure XP beads (Beckman Coulter, Inc., Brea, CA, USA). The quantified library pool was subsequently delivered to Novogene (Novogene Corporation Inc., Sacramento, CA, USA) for 2×150 bp paired-end sequencing on the Illumina NovaSeq X Plus platform (Illumina, Inc., San Diego, CA, USA).

### 2.6 Taxonomic Classification of and Functional Annotation Metagenomic Sequences

The raw FASTQ files were first assessed for quality using FastQC (v0.11.6)^13^ Adapter removal and quality trimming were then performed with Trimmomatic (v0.39)^14^, applying stringent criteria: a 4-base sliding window, a minimum Phred quality threshold of 20, and a minimum retained read length of 50 bp. Following initial quality filtering, an average of 51.84 million read pairs per sample (range: 22.30–77.72 million) were retained for subsequent analysis. To isolate microbial sequences, host-derived reads were removed in silico using BBduk (BBtools package v38.x)^15^. This was achieved by aligning the quality-trimmed reads to the Gallus gallus reference genome (assembly GRCg6a, accession GCA_000002315.5) with a k-mer length of 31; reads that did not align were classified as “host-filtered” and used for all subsequent analyses. Host read filtering effectively removed a small proportion of host-derived sequences across all samples, with the majority of reads (94–99%) retained as unmapped for downstream microbial analysis (Data S1). Taxonomic profiling was performed on the host-filtered reads using Kraken2 (v2.1.x) with a custom database (latest update July 2025). The classification was run with a confidence threshold of 0.1 and a requirement for a minimum of 2 hit groups. To refine the abundance estimates at the species level, the Kraken2^16^ output was processed with Bracken (Bayesian Reestimation of Abundance after Classification with KrakEN)^17^. Finally, taxonomic identifiers were mapped to their full scientific nomenclature using TaxonKit^18^. Functional profiling of antimicrobial resistance (AMR) and virulence-associated genes was conducted using the AMR++ pipeline^19^. High-quality reads were aligned to the MEGARes database (v2.0)^19^ and the experimentally verified Set-A from the Virulence Factor Database (VFDB, August 2024)^20^ (with BWA^21^. To maintain specificity, only gene hits with at least 99% sequence coverage (i.e., reads covering ≥99% of the gene length) were retained for downstream analysis. The resulting high-confidence gene assignments were further processed with ResistomeAnalyzer (v2.1; https://github.com/cdeanj/resistomeanalyzer) to profile antimicrobial resistance patterns. This rigorous filtering step effectively reduced partial alignments and improved the accuracy of resistome characterization. Reads Per Kilobase (RPK) gene family abundance tables obtained from HUMAnN3^22^ were refined by merging duplicate UniRef90 identifiers^23^ and removing associated taxonomic labels. To eliminate redundancy, gene abundances corresponding to each UniRef90 ID were summed. KEGG^24^ Orthology (KO) identifiers were then assigned using the predefined mapping file map_ko_uniref90.txt, which links UniRef90 IDs to their respective KEGG Orthologs. For genes mapped to multiple KOs, abundances were proportionally divided among all associated KOs. Individual KO abundance tables were generated for each sample, presenting normalized counts per KEGG Ortholog. These tables were subsequently combined into a single comprehensive matrix, with absent KOs replaced by zeros to maintain uniformity. Functional annotation was conducted using the KEGGREST R package^25^ to associate KOs with KEGG pathways, facilitating pathway-level abundance profiling and visualization. This analytical framework provided detailed metabolic insights while ensuring compatibility with subsequent statistical and comparative analyses.

### 2.6 Statistical and Bioinformatic Analysis

Microbial diversity analyses were conducted in R (v4.2.0) using RStudio (2022.07.1), primarily leveraging the phyloseq (v1.50.0)^26^ and vegan (v2.6-4)^27^ packages. Alpha diversity was evaluated using observed taxon richness and the Shannon diversity index, calculated from total sum scaling (TSS)-normalized data. To assess differences between groups, non-parametric tests were applied: Wilcoxon rank-sum tests for two-group comparisons, and Kruskal-Wallis tests followed by Dunn’s post hoc tests with Bonferroni correction for multi-group comparisons. Beta diversity was quantified using Bray-Curtis dissimilarity matrices^28^ derived from TSS-normalized counts. Permutational multivariate analysis of variance (PERMANOVA) was performed using the adonis2 function with 999 permutations to evaluate associations between microbial community composition and clinical variables such as infection type, gender, and comorbidities, with R² values reported as effect sizes. Community dissimilarity patterns were visualized using principal coordinates analysis (PCoA) via phyloseq’s ordinate function. All analyses accounted for the compositional nature of microbiome data, and false discovery rate was controlled using the Benjamini-Hochberg procedure, with q-values < 0.05 considered statistically significant. Associations among the top 50 most abundant microbial species (based on relative abundance) were assessed using Spearman’s correlation, with strong correlations (|r| > 0.6) visualized as chord diagrams using the circlize package^29^. Additionally, relationships between bacterial species, antimicrobial resistance classes and virulence factors were explored using Spearman’s rank correlation via the Hmisc^30^ and corrplot^31^ packages. Data normality was evaluated with the Shapiro-Wilk test, and non-parametric tests were justified due to the predominance of non-normal distributions (p < 0.05). All relative abundance values were derived from shotgun metagenomic sequencing, with statistical significance set at α = 0.05.

## 3. Results

### 3.1 Effects of Spent Lees and Autoclaved Spent Lees on Body Weight in Poultry

The impact of dietary supplementation with spent Lees and autoclaved spent Lees on body weight gain in poultry was evaluated over a 14-day period (Table 2). At baseline (Day 0), no significant differences were observed among the control (98.4 ± 5.0 g), spent Lees (86.2 ± 11.4 g), and autoclaved spent Lees (90.2 ± 5.1 g) groups, confirming uniform starting weights across treatments. By Day 7, both spent Lees-supplemented groups exhibited marked increases in body weight relative to the control. The spent Lees group showed the highest average weight (251.4 ± 21.4 g), followed by the autoclaved spent Lees group (239.6 ± 11.4 g), whereas the control group averaged 213.4 ± 6.1 g. These results indicate that spent Lees supplementation enhanced early growth performance, with spent Lees producing a more pronounced effect. At Day 14, the growth-promoting effects of spent Lees supplementation persisted. The spent Lees group achieved an average body weight of 545.4 ± 66.3 g, while the autoclaved spent Lees group reached 573.4 ± 17.2 g, both exceeding the control group (526.2 ± 18.6 g). Interestingly, although spent Lees accelerated early growth, birds receiving autoclaved spent Lees ultimately achieved higher final weights, suggesting potential differences in growth kinetics between the two spent Lees preparations. Supplementation with either spent Lees or autoclaved spent Lees significantly improved body weight gain compared to the control, highlighting the potential of spent Lees as a dietary growth promoter in poultry.

### 3.2 Spent Lees Supplementation Alters the Gut Microbiota Structure in Chickens

To assess the impact of dietary spent Lees on the ecological structure of the chicken gut microbiota, we first evaluated within-sample (alpha) diversity. The Control group at day 0 exhibited a baseline Shannon diversity index of 2.30. By day 14, the Control group maintained a similarly high alpha diversity (Shannon index: 2.28 - 2.34). In contrast, both spent Lees-supplemented groups showed a marked reduction in diversity at day 14. The Autoclaved Spent Lees group had intermediate values (Shannon index: 1.54 - 1.96), while the Spent Lees group displayed the lowest diversity (Shannon index: 1.41 - 1.61) (Fig. 1A, Data S1). We next investigated the differences in microbial community composition between groups using Bray-Curtis dissimilarity. Permutational multivariate analysis of variance (PERMANOVA) revealed that the dietary treatment group was a significant factor explaining the variance in the gut microbiota structure (adonis2, R² = 0.989, F = 91.93, p = 0.012). This indicates a strong and statistically significant separation of the microbial communities based on whether the chickens received the control diet, spent Lees, or autoclaved spent Lees (Fig 1B).

**Fig. 1:**
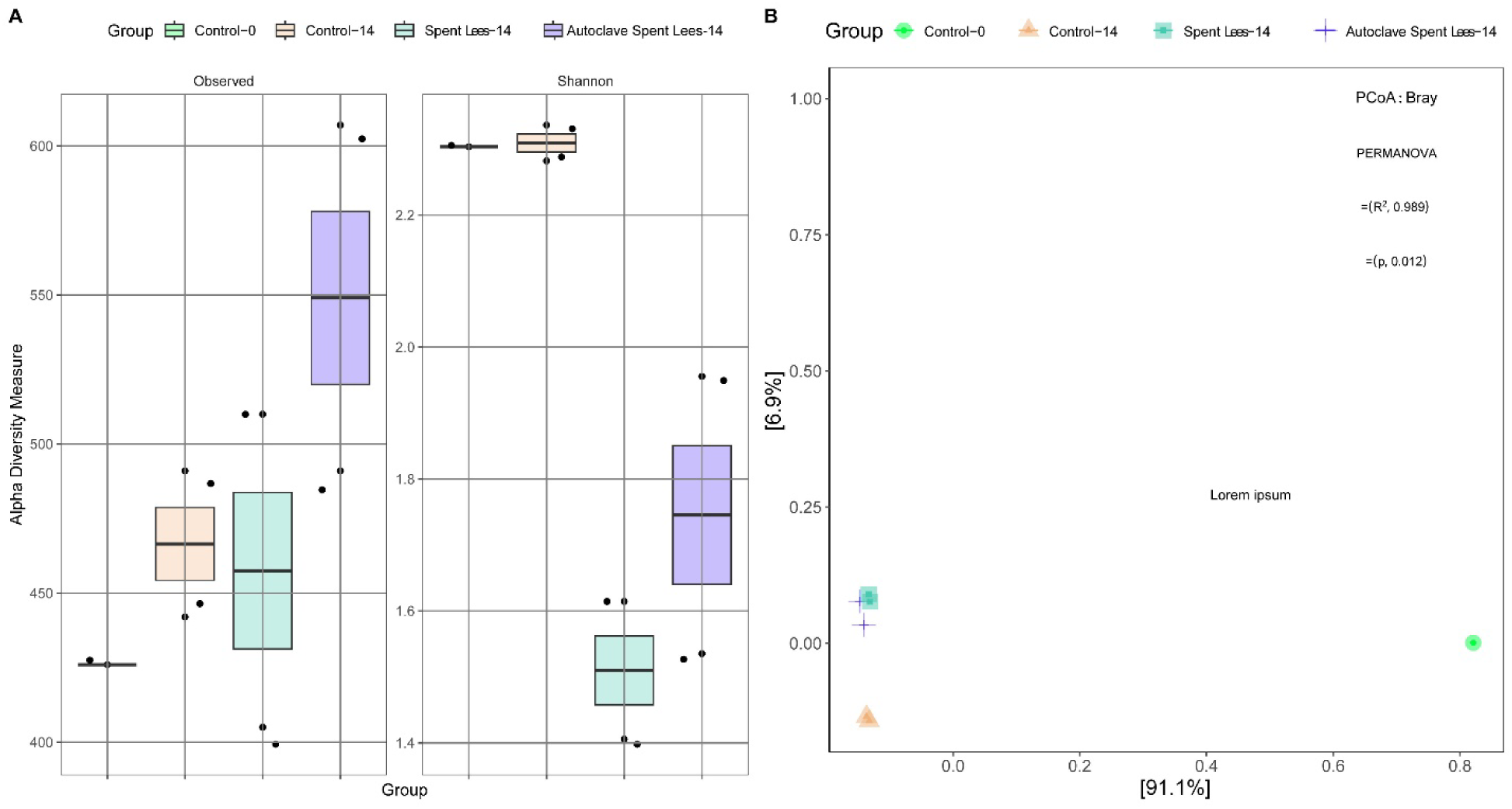
Assessment of microbial diversity within and between groups. (A) Within-sample (alpha) diversity was measured with the Observed Richness and Shannon Diversity Index. The boxplots illustrate the distribution of these metrics across groups, with individual data points shown. (B) Between-sample (beta) diversity, representing community composition differences, was visualized using a Principal Coordinates Analysis (PCoA) plot based on Bray-Curtis dissimilarity. PERMANOVA with 999 permutations was used to test the statistical significance of the groupings by group and time frame.

### 3.3 Dietary Spent Lees Drives a Major Phylogenetic Restructuring of the Gut Microbiome

Analysis of the microbial community composition revealed profound shifts at the phylogenetic level in response to dietary supplementation. The gut microbiota was overwhelmingly dominated by the domain Bacteria, which constituted >99.9% of the community in all spent Lees-supplemented groups and 99.3% in the Control-14 group (Fig. 2A, Data S1). A notable finding was the significant reduction of eukaryotic DNA in the spent Lees-fed groups (0.01% in Spent Lees and Autoclaved Spent Lees) compared to the Control-0 group (0.62%), suggesting that dietary spent Lees effectively outcompetes or suppresses other gut eukaryotes. At the phylum level, a dramatic restructuring was observed (Fig. 2B). The baseline (Control-0) microbiota was co-dominated by Bacillota (53.3%) and Pseudomonadota (45.0%), with low levels of Bacteroidota (0.6%). By day 14, the Control group exhibited an expected developmental shift, characterized by a decrease in Bacillota (40.4%) and Pseudomonadota (0.9%), and a substantial expansion of Bacteroidota (57.3%). However, spent Lees supplementation profoundly amplified this shift towards Bacteroidota. The Autoclaved Spent Lees group was dominated by Bacteroidota (69.9%) and Bacillota (27.3%), while the Spent Lees group showed an even more extreme phenotype, with Bacteroidota soaring to 75.5% and Bacillota suppressed to 21.8%. Concurrently, the relative abundance of Actinomycetota was higher in all day-14 groups compared to baseline, with the Autoclaved Spent Lees group showing the highest level (1.5%). These results demonstrate that both spent Lees and autoclaved spent Lees drive the gut microbiome towards a Bacteroidota-dominated state, with spent Lees exerting a more potent effect.

**Fig. 2:**
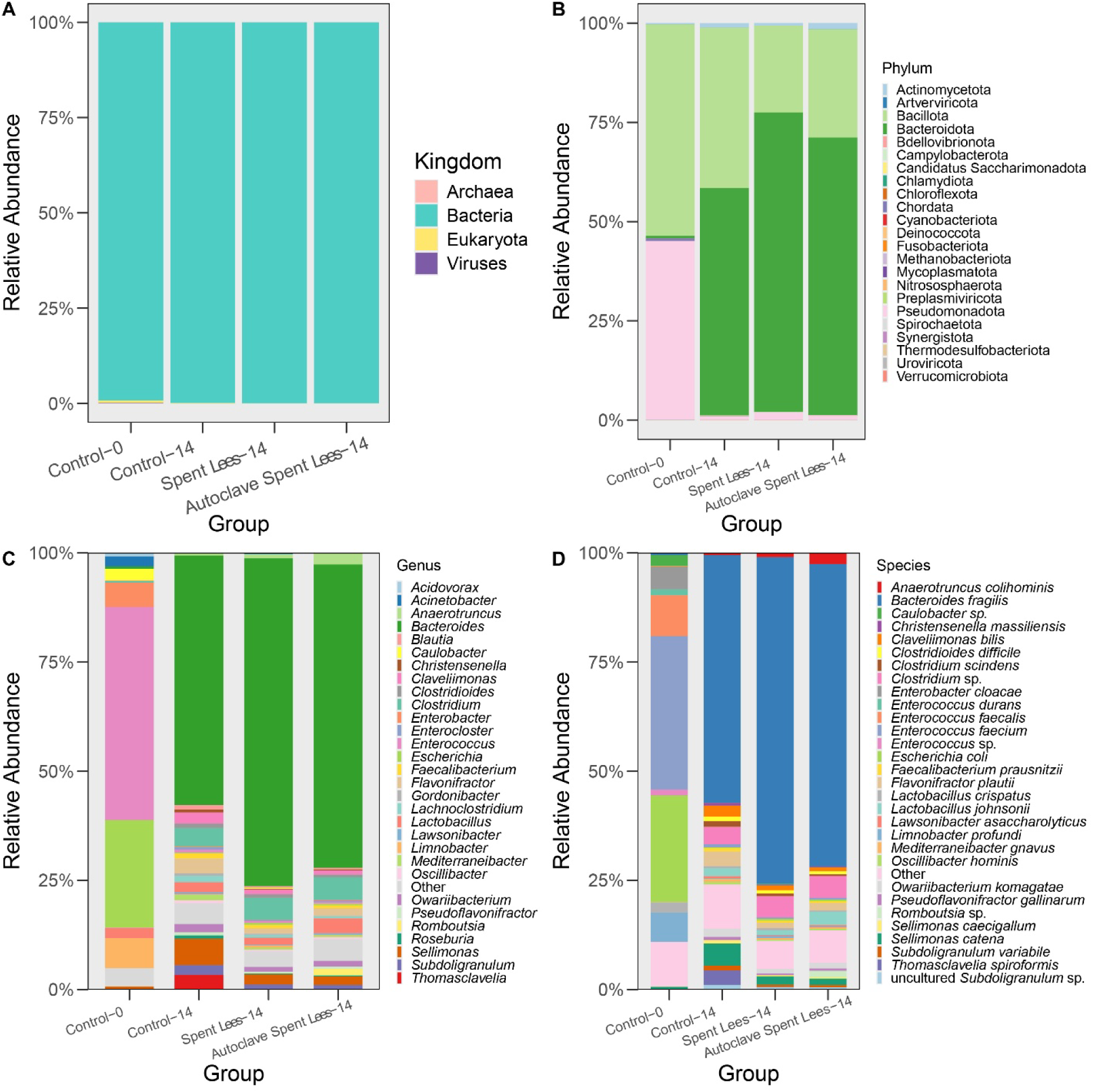
Stacked bar plots depicting the taxonomic composition of microbial communities across samples. The relative abundance is shown at different taxonomic levels: (A) Kingdoms, (B) Phyla, (C) the top 30 Genera, and (D) the top 30 most abundant Species. Samples on the x-axis are grouped and ordered alphabetically. The y-axis represents the relative abundance (%), and each taxon is indicated by a distinct color.

### 3.4 Spent Lees Supplementation Drives a Genus-Level Microbial Shift Characterized by Bacteroides Dominance and Pathogen Exclusion

Building on the profound phylum-level changes, we investigated the specific genera modulated by dietary spent Lees. The most striking effect was the dramatic enrichment of the genus Bacteroides. While it constituted a mere 0.53% of the baseline (Control-0) community, its abundance expanded to 57.17% in the Control-14 group, reflecting normal maturation. However, spent Lees supplementation supercharged this expansion, with Bacteroides reaching 69.45% in the Autoclaved Spent Lees group and a remarkable 75.09% in the Spent Lees group, establishing it as the dominant taxon (Fig. 2C, Data S1). Concurrently, spent Lees supplementation led to the near-complete exclusion of potentially pathogenic or opportunistic genera that were prominent in the early-life microbiome. Most notably, the high initial abundances of Enterococcus (48.81% in Control-0) and Escherichia (24.57% in Control-0) were drastically reduced in all day-14 groups. However, this suppression was significantly more pronounced in the spent Lees-fed groups, where their combined relative abundance was kept below 1%, compared to nearly 1% in the Control-14 group. Other environmental or opportunistic genera, such as *Acinetobacter*, *Caulobacter*, and *Limnobacter*, which were abundant at baseline, were also effectively suppressed to negligible levels (<0.5%) in the spent Lees-treated chickens. Beyond the dominance of Bacteroides, the spent Lees-supplemented groups exhibited a consistent modification of other commensal and fermentative genera. Several short-chain fatty acid (SCFA)-producing genera within the *Bacillota* phylum, including *Blautia*, *Faecalibacterium*, *Mediterraneibacter*, *Subdoligranulum*, and *Roseburia*, showed a graded response: their abundances were generally highest in the Control-14 group, intermediate in the Autoclaved Spent Lees group, and lowest in the Spent Lees group. This suggests that the extreme proliferation of Bacteroides in the Spent Lees group may come at the expense of a more diverse SCFA-producing consortium. Furthermore, the Spent Lees group displayed a specific and significant reduction in the genus Lactobacillus compared to all other groups. A notable finding was the differential response of the genus *Thomasclavelia* (formerly *Clostridium* cluster XI), which was highly abundant in the Control-14 group (3.36%) but was substantially lower in both the Autoclaved (0.03%) and Spent Lees (0.08%) groups, indicating a potent suppressive effect of spent Lees on this specific clostridial taxon. Dietary spent Lees, particularly in its live form, orchestrates a gut microbial community characterized by extreme dominance of Bacteroides, a profound exclusion of early-life enteropathogens, and a restructuring of the commensal fermentative community.

### 3.5 Spent Lees Supplementation Eradicates Enteropathogens and Drives a Monodominant Gut Ecosystem by a Single Bacteroides fragilis Strain

Our deep sequencing revealed that the microbial restructuring induced by spent Lees supplementation was extraordinarily specific, culminating in a dramatic alteration of the gut microbial landscape at the species level. The most profound effect was the near-total eradication of opportunistic and pathogenic bacteria that dominated the early-life (Control-0) microbiome. Species including Enterococcus faecium (35.18% in Control-0), Escherichia coli (24.45%), and Enterobacter cloacae (5.21%) were suppressed to negligible abundances (<1%) in all spent Lees-supplemented groups, an effect that was significantly more pronounced than in the aged Control-14 group (Fig. 2D, Data S1). Concurrently, we observed a remarkable, strain-level dominance by a single species: Bacteroides fragilis. While the baseline community contained a low abundance of B. fragilis (0.53%), it underwent a natural expansion in the Control-14 group to 56.70% (Fig. 3). However, dietary spent Lees supercharged this expansion. In the Autoclaved Spent Lees group, B. fragilis reached 69.23%, and in the Spent Lees group, it achieved a striking 74.83% relative abundance, effectively becoming the monodominant species in the gut. This suggests that the spent Lees-derived nutrients create a powerful selective advantage specifically for B. fragilis over all other microbial taxa. The proliferation of B. fragilis occurred alongside a significant reshaping of the commensal Clostridia. Several beneficial, short-chain fatty acid (SCFA)-producing species, such as *Faecalibacterium prausnitzii*, *Subdoligranulum variabile*, and *Blautia hydrogenotrophica*, were most abundant in the Control-14 group, but their levels were progressively reduced in the Autoclaved and Spent Lees groups, indicating that the extreme fitness of B. fragilis may partially outcompete these commensals. Furthermore, the Spent Lees group exhibited a specific and significant reduction in the beneficial species *Lactobacillus johnsonii* (1.06% in Spent Lees vs. 1.86% in Control-14 and 3.08% in Autoclaved Spent Lees). A notable and potentially beneficial exclusion was observed for the species *Thomasclavelia spiroformis*, which reached 3.27% in the Control-14 group but was suppressed to 0.05% and 0.08% in the Spent Lees and Autoclaved Spent Lees groups, respectively. This indicates a potent inhibitory effect of spent Lees on this specific, and often undesirable, clostridial species. Spent Lees supplementation does not merely modulate the gut microbiome; it orchestrates a dramatic ecological shift characterized by the eradication of key enteropathogens and the ascendance of Bacteroides fragilis to a position of monodominance, fundamentally restructuring the gut ecosystem.

**Fig. 3:**
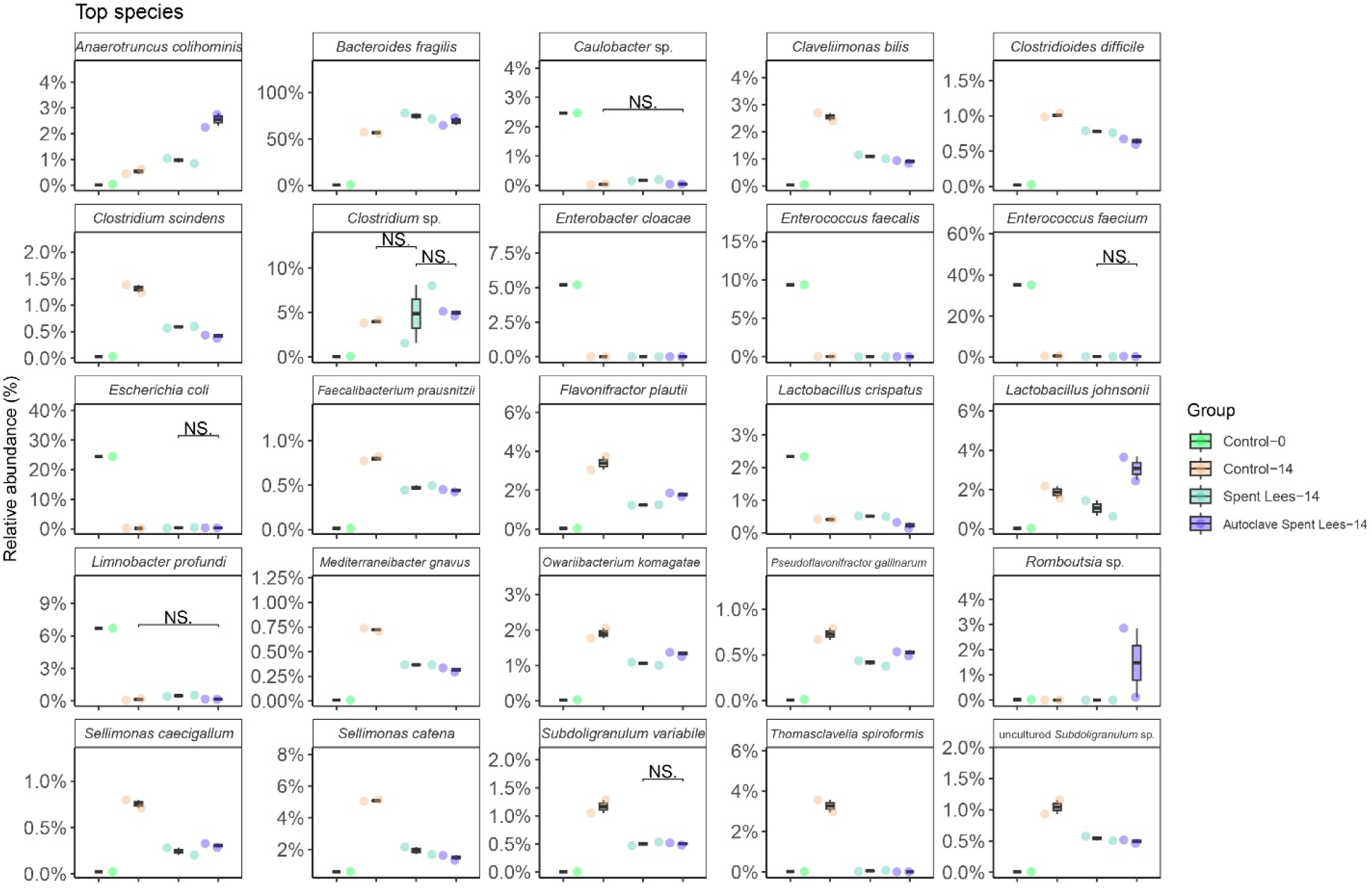
Box-plot analysis of the top 25 species across samples. Box plots were generated to visualize the distribution of the top 50 microbial species across different sample groups. Statistical comparisons were performed using the Wilcoxon rank-sum test, where significance levels are indicated as *p < 0.05, **p < 0.01, and ***p < 0.001.

### 3.6 Gut Virome Depletion is Linked to Loss of Bacterial Hosts During Development and Spent Lees Supplementation

Analysis of viral reads revealed that the early-life (Control-0) chicken gut microbiome contained a detectable population of bacteriophages, primarily from the Uroviricota phylum (0.11% of total reads). Notably, these viral sequences were dominated by members of the Copernicusvirus genus, which includes bacteriophages known to target enterococci. This viral signature was entirely absent in all day-14 groups, regardless of diet. No detectable levels of Uroviricota, Copernicusvirus, or any other viral taxa were present in the Control-14, Spent Lees-14, or Autoclaved Spent Lees-14 groups (Data S1). The disappearance of these phages is temporally linked to the dramatic reduction in their putative bacterial hosts. For instance, Enterococcus faecium—a primary host for Copernicusvirus—collapsed from 35.18% abundance at Day 0 to below 0.63% in all day-14 groups (Data S1). These findings indicate that the depletion of the initial gut virome is a common outcome of post-hatch development under these husbandry conditions, driven by the loss of specific host bacteria. While spent Lees supplementation was a powerful driver of this bacterial shift, the resulting virome depletion was consistent with the broader pattern observed in the aged control group.

### 3.7 Network Analysis Reveals a Spent Lees-Driven Bifurcation of the Gut Microbiome into Two Distinct Ecological Consortia

To move beyond taxon-specific shifts and understand the system-level reorganization of the microbial community, we constructed a co-occurrence network based on Spearman correlations among the top 50 most abundant species. This analysis revealed that the gut microbiome is structured into two distinct, anti-correlated ecological consortia (Fig. 4, Data S1). The first consortium, which we term the “;Mature Commensal Network,” is a tightly correlated group of strictly anaerobic, fermentative bacteria that became dominant in the day-14 groups. This network was characterized by exceptionally strong positive correlations (Spearman’s ρ > 0.9) between key species such as *Clostridium scindens, Blautia hydrogenotrophica, Sellimonas catena, Clostridioides difficile*, and *Ruminococcus torques*. This consortium also included well-known butyrate producers like *Faecalibacterium prausnitzii* and *Subdoligranulum variabile,* which co-occurred strongly with the group (ρ = 0.96 and 0.86 with *C. scindens*, respectively). The dominance of Bacteroides fragilis was integrated into this network, showing strong positive correlations with other Bacteroides species like *B. thetaiotaomicron* (ρ = 0.93 with *C. scindens*), positioning it as a keystone member of this mature community structure. Strikingly, this entire Mature Commensal Network was strongly and universally anti-correlated with a second consortium, the “Early-Life and Environmental Network.” This group was dominated by host-associated and environmental taxa that were abundant at Day 0, including *Enterococcus faecium*, and *Escherichia coli* reads. This negative relationship was stark; for example, *Bacteroides fragilis* was strongly anti-correlated with E. faecium (ρ = −0.89) and E. coli (ρ = −0.86 with *B. thetaiotaomicron*). The transition from the early-life to the mature state, whether driven by age or dramatically accelerated by spent Lees supplementation, is therefore not a series of independent changes but a coordinated ecological shift. The data illustrate a fundamental biological trade-off: the proliferation of a synergistic network of obligate anaerobes necessitates the exclusion of a separate, facultative anaerobic consortium associated with the immature gut and external environment.

**Fig. 4:**
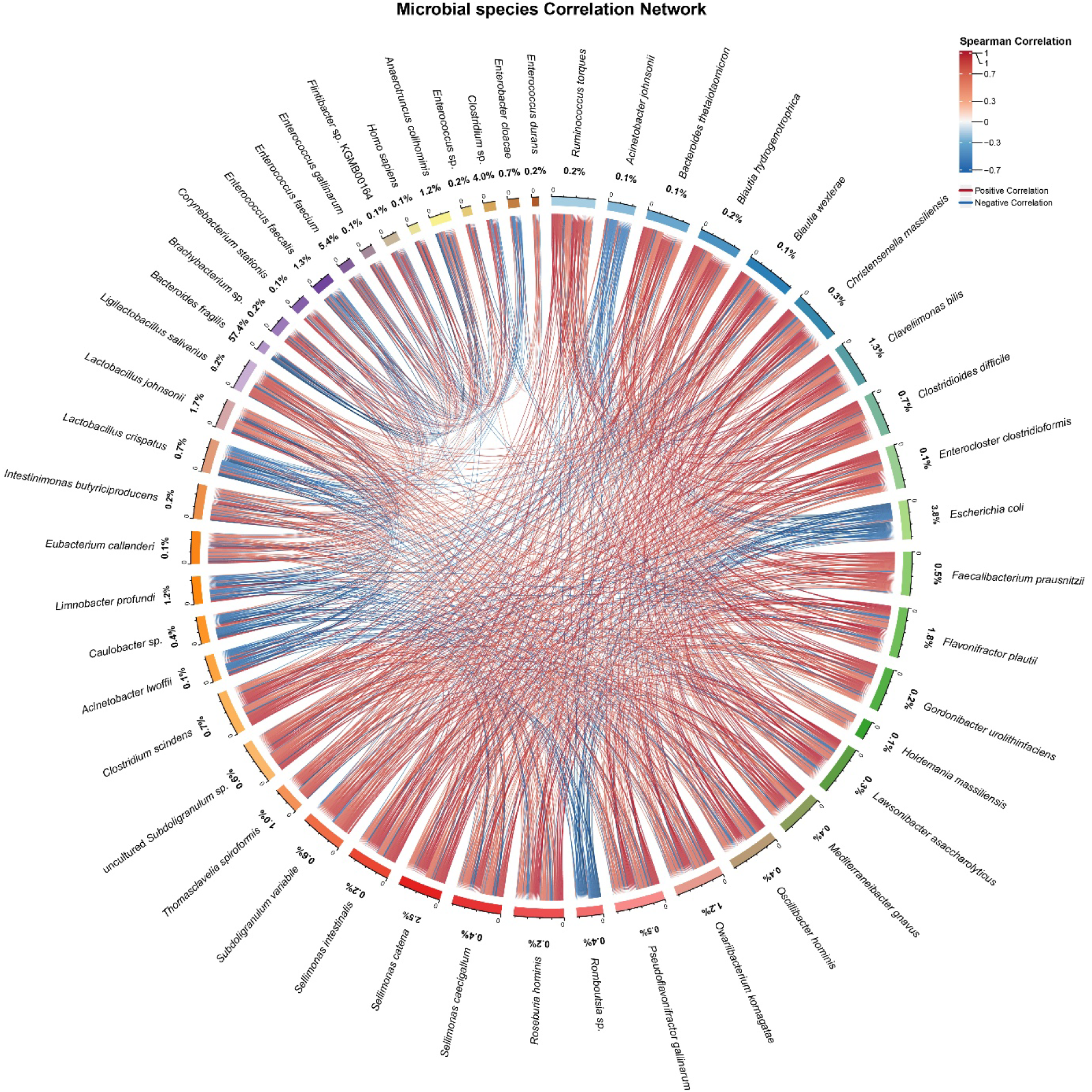
Network analysis of microbial co-occurrence. Interactions were inferred from Spearman correlation analysis of the 50 most prevalent species. The chord diagram visualizes statistically significant, robust correlations (|r| > 0.6). Band dimensions and color saturation are proportional to the correlation strength, with orange (positive) and moss green (negative) distinguishing the nature of the relationship. Visualization was performed with the *circlize* package.

### 3.8 Spent Lees Supplementation Mitigates the Natural Expansion of the Gut Resistome

Given the profound restructuring of the gut microbiome, we investigated its functional consequences by profiling the abundance of antibiotic resistance genes (ARGs), collectively known as the “resistome.” We observed a dramatic and natural expansion of the resistome in the gut of chickens as they aged, which was significantly modulated by dietary spent Lees supplementation. The baseline (Control-0) resistome was characterized by a high abundance of genes conferring resistance to biocides, metals, and multi-drug efflux pumps, consistent with an early-life microbiome exposed to environmental challenges (Data S2). However, by day 14, the control-fed chickens (Control-14) exhibited a massive proliferation of specific, high-risk ARGs.

Most notably, genes for tetracycline ribosomal protection (e.g., tetW, 18.35%) and oxazolidinone resistance (o23S, 10.83%) became dominant. Concurrently, a suite of genes for aminoglycoside modification (e.g., ant6, 9.00%; aph3’’-IIIa, 5.26%), MLS resistance (e.g., cfr, 5.96%), and other drug classes expanded markedly, creating a highly multi-drug resistant gut environment. Strikingly, spent Lees supplementation (Spent Lees-14) substantially curtailed this resistome expansion. The abundance of the dominant tetW gene was reduced from 18.35% to 15.48%. More profound suppressive effects were observed for other key genes; the oxazolidinone resistance gene o23S was undetectable in the Spent Lees group (compared to 10.83% in Control-14), and the lincosamide resistance gene lnuC was reduced from 1.70% to 1.51%. A similar trend of reduction was observed across multiple drug classes in the autoclaved spent Lees group, though the effects were generally less pronounced than with spent Lees. This reshaping of the resistome was not limited to antibiotics. Spent Lees supplementation also led to a significant reduction in the abundance of genes conferring resistance to biocides (e.g., acid resistance genes gadA, gadC) and metals (e.g., multi-metal efflux pumps cusA, cusB), which were highly abundant in the Control-0 group. This suggests that spent Lees induces a global attenuation of the microbial stress response arsenal. The maturation of the chicken gut under standard conditions is associated with a concerning bloom of diverse and high-abundance ARGs. Dietary supplementation with spent Lees presents a compelling strategy to mitigate this expansion, significantly reducing the abundance of critically important resistance genes and potentially lowering the risk of disseminating antimicrobial resistance from poultry production.

To elucidate the drivers behind the profound resistome restructuring, we performed a correlation network analysis between the top 50 microbial species and all detected antimicrobial resistance gene (ARG) classes. This analysis revealed that the resistome is not a random assemblage but is intrinsically linked to the two distinct ecological consortia we identified earlier, with ARGs acting as biomarkers for these community states. We found that the “Early-Life and Environmental Network” was a major reservoir for a broad spectrum of resistance genes. Species characteristic of this consortium, such as Escherichia coli, showed strong positive correlations with ARGs conferring resistance to fluoroquinolones, aminocoumarins, multi-drug efflux pumps, and biocides (e.g., acid resistance genes gadA/gadC, ρ > 0.85, p < 0.05) (Fig. 5, Data S2). This suggests that the early-life gut and its associated environmental bacteria are intrinsically equipped with a diverse defensive arsenal against chemical stressors. Conversely, the “Mature Commensal Network” was defined by a contrasting resistome profile. Key members of this consortium, including *Blautia hydrogenotrophica, Clostridioides difficile*, and *Sellimonas catena*, were strongly anti-correlated with the vast majority of biocide and multi-drug efflux resistance genes (ρ < −0.78, p < 0.05). Instead, their abundance was positively correlated with a specific set of ARGs that became dominant in the day-14 groups, most notably genes for tetracycline ribosomal protection (tetW, tetO, tetQ) and MLS resistance (cfr, erm genes). This pattern reveals a fundamental ecological shift: spent Lees-driven maturation replaces a community equipped for generic environmental stress resistance with one specialized for intra-microbial competition. The dominance of Bacteroides fragilis and the associated mature network correlates with a resistome focused on outcompeting other bacteria (e.g., via tetracycline and macrolide resistance mechanisms), rather than surviving host-derived or external chemical assaults. The reduction in high-risk ARGs observed with spent Lees supplementation is a direct consequence of a trophic shift in the gut ecosystem. By driving the exclusion of the early-life, environmentally-adapted resistome reservoir and promoting a stable, commensal-dominated community, spent Lees effectively redirects the functional capacity of the microbiome away from a broad-spectrum defensive state.

**Fig. 5:**
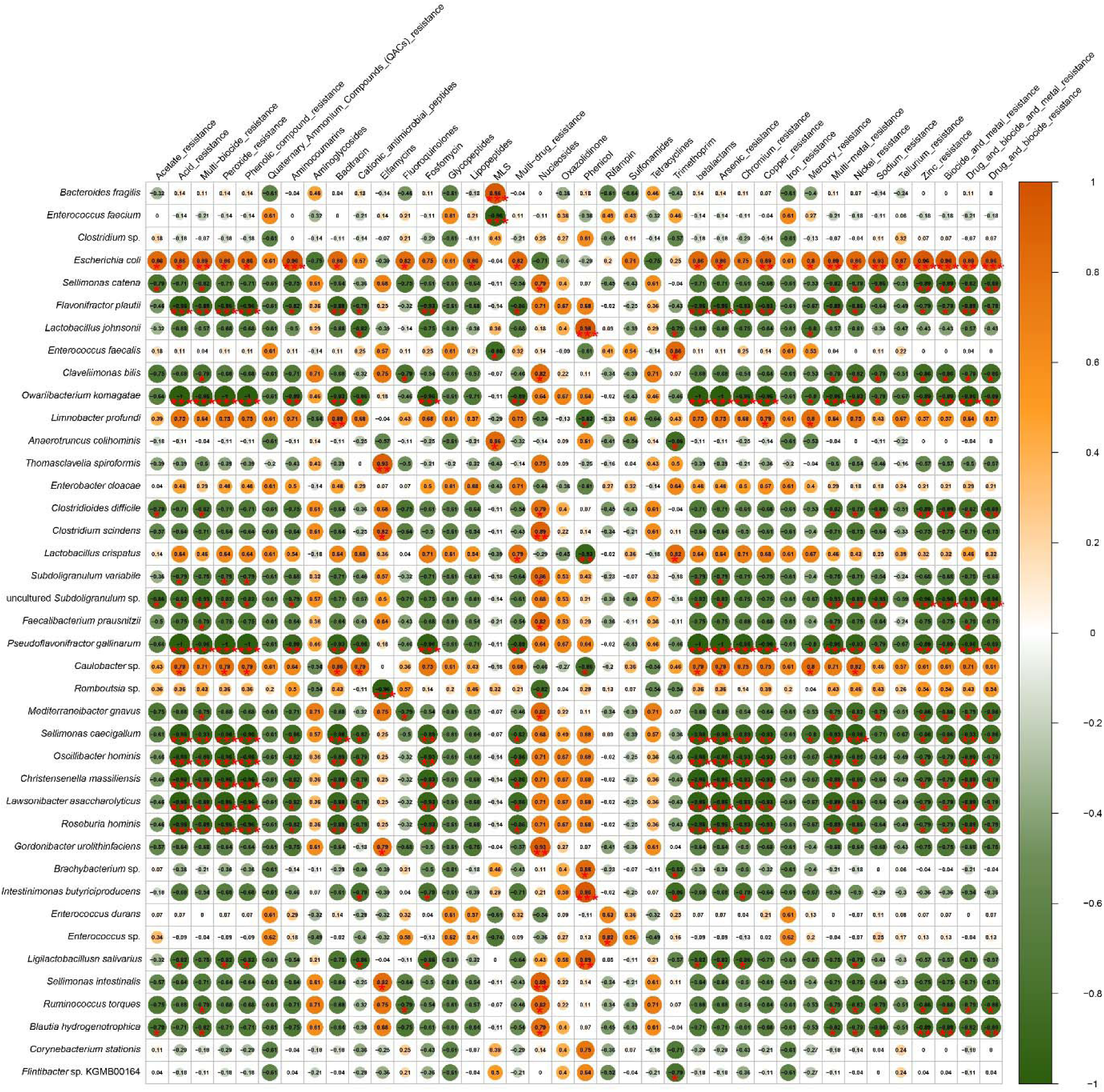
Association of dominant bacterial species with antibiotic resistance classes. A correlation heatmap displays Spearman’s rank coefficients between the relative abundance of the top 50 most abundant bacterial species (y-axis) and the abundance of various antibiotic resistance classes (x-axis). Statistically significant correlations are represented by circles, with color (orange for positive, green for negative) and size scaled to the magnitude of the correlation coefficient. Significance levels are indicated as *p < 0.05, **p < 0.01, and ***p < 0.001.

### 3.9 Spent Lees Supplementation Restructures the Gut Virulome by Suppressing Pathogenic Effector Systems

To assess the potential impact of dietary spent Lees on the pathogenic potential of the gut microbiome, we profiled the abundance of virulence factor genes (VFGs), collectively termed the “virulome.” We observed that spent Lees supplementation induced a significant restructuring of the virulome, characterized by the suppression of high-impact virulence mechanisms and a shift towards commensal-associated colonization factors. The baseline (Control-0) and Control-14 gut microbiomes were enriched with genes encoding sophisticated Type III Secretion System (T3SS) effectors, which are critical for host cell invasion and pathogenesis by enteric pathogens. Genes such as espX4 (1.48% in Control-0, 3.38% in Control-14) and espL4 (1.59% in Control-0, 1.96% in Control-14) were prominent. In contrast, both spent Lees and autoclaved spent Lees supplementation markedly reduced the abundance of these effector genes (e.g., espX4: 2.50% in Spent Lees; espX1: 0.63% in Spent Lees vs. 1.00% in Control-0) (Data S2). Concurrently, spent Lees supplementation enriched for genes associated with iron acquisition and basic adherence. Genes involved in enterobactin synthesis (e.g., entE) and hemin transport (e.g., shuV, iroC) were maintained or increased in spent Lees-fed groups. Similarly, genes for Type 1 fimbriae assembly (e.g., fimC, fimD, fimH), which facilitate commensal colonization, were consistently present across all groups but showed a stable or slightly elevated profile in spent Lees-supplemented chickens. This shift reflects a change in microbial strategy: spent Lees promotes a virulome dominated by factors for nutrient scavenging and mucosal persistence, while suppressing the expression of the molecular weaponry, the T3SS effectors, used for host cell subversion and disease. This suggests that spent Lees fosters a community where bacteria are optimized for peaceful coexistence within the gut niche rather than aggressive host invasion. Spent Lees supplementation does not eliminate virulence capacity but fundamentally redirects it, favoring a commensal lifestyle over an overtly pathogenic one. This reprogramming of the gut virulome towards colonization factors and away from offensive effector systems represents a key mechanism by which spent Lees may enhance gut barrier integrity and overall host well-being.

To define the relationship between microbial identity and pathogenic potential, we integrated the abundance of the top 50 species with virulence factor gene (VFG) classes. This analysis revealed that the two previously identified ecological consortia are defined not only by their taxonomy and resistome but also by distinct virulome signatures, providing a unified functional genomic profile for each community state. The “Mature Commensal Network”—comprising species such as *Blautia hydrogenotrophica, Clostridioides difficile*, and *Sellimonas catena*—was strongly and consistently associated with VFGs for Adherence (e.g., fim genes; ρ > 0.78, p < 0.05) and global Regulation (e.g., rpoS; ρ > 0.85, p < 0.05) (Fig. 6, Data S2). This correlation profile underscores a strategy of persistent mucosal colonization and environmental stress adaptation, hallmarks of a stable, commensal lifestyle focused on niche persistence. Conversely, the “Early-Life and Environmental Network” was characterized by an inverse relationship with these communal traits. More notably, the dominance of the keystone commensal *Bacteroides fragilis* was strongly anti-correlated with the presence of classic pathogens like *Enterococcus faecium* and *Enterococcus faecalis* (ρ < −0.78, p < 0.05), which themselves are known to harbor diverse virulence factors. Furthermore, we observed a significant decoupling between different virulence strategies. Genes for the Effector Delivery System (e.g., T3SS) were strongly anti-correlated with global Regulation (ρ = −0.857, p = 0.014), suggesting a fundamental trade-off between the energy-intensive maintenance of offensive molecular weaponry and the regulatory networks that facilitate a commensal, stress-adapted existence. The spent Lees-driven transition to a Bacteroides-dominated ecosystem enforces a functional outcome where the virulome is canalized towards traits supporting commensalism. The strong correlations between specific bacterial consortia and discrete VFG classes demonstrate that virulence potential is an emergent property of the community structure, which can be steered towards a less pathogenic state through dietary intervention.

**Fig. 6:**
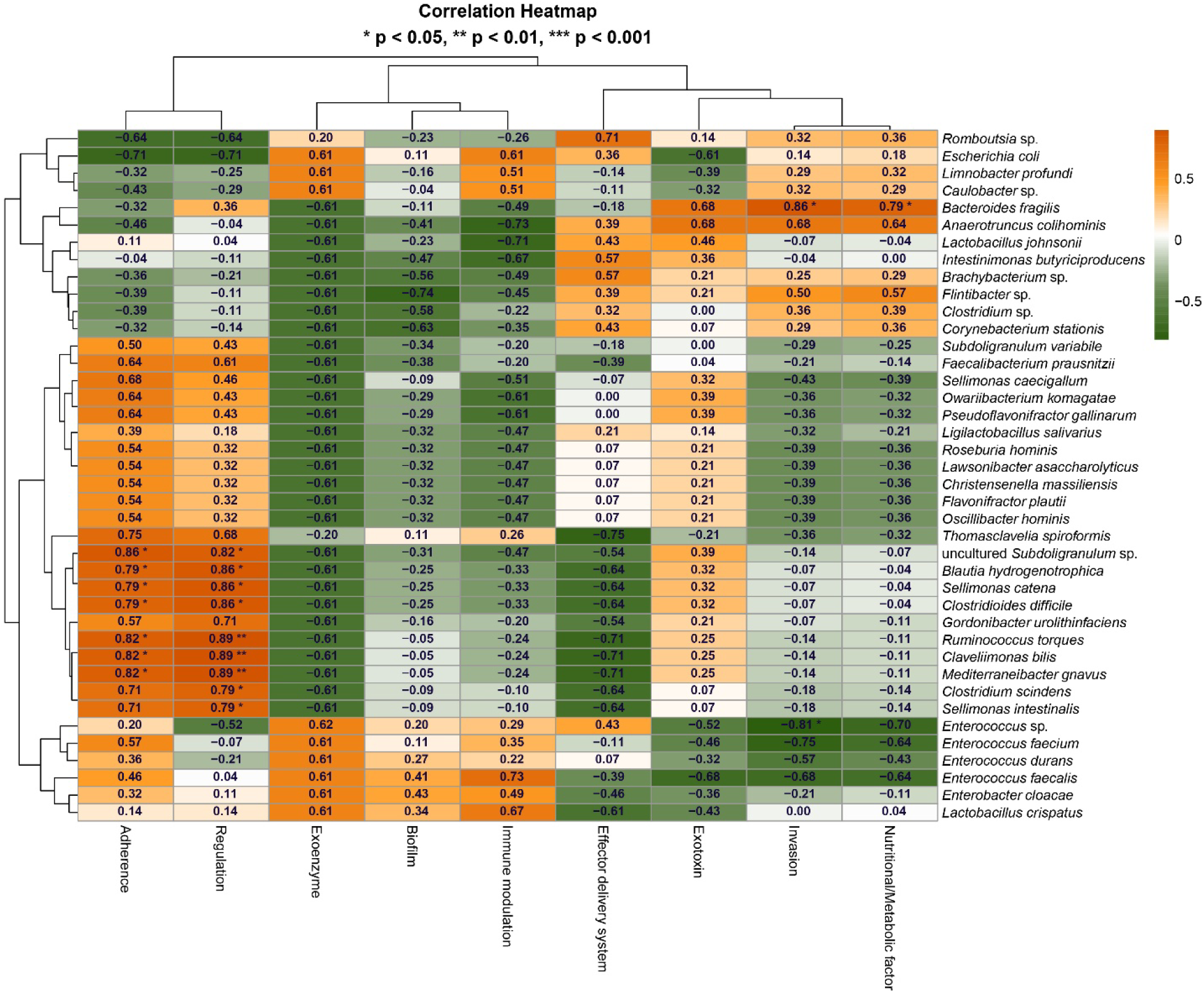
Spearman’s correlation matrix between Virulence Factors and the top 50 dominant bacterial species. The heatmap displays Spearman’s correlation coefficients (r) between measured Virulence Factors (X-axis) and the relative abundance of the top 50 most abundant bacterial species (Y-axis). Hierarchical clustering dendrograms are shown for both axes. Positive and negative correlations are represented by vibrant orange and moss green rectangles, respectively. The color intensity of each rectangle are proportional to the strength of the correlation (|r|). Statistical significance is denoted as: *p < 0.05, **p < 0.01, ***p < 0.001.

### 3.10 Spent Lees Supplementation Reprograms the Gut Microbiome for Enhanced Biosynthesis and Host-Microbe Symbiosis

To understand the systemic functional consequences of the spent Lees-induced microbial shift, we analyzed the gut metagenome using the KEGG database. This revealed a profound reprogramming of the community’s metabolic potential, moving from an environmentally-focused state to one optimized for host-associated biosynthesis and nutrient exchange. The gut microbiome of control-fed chickens at day 0 (Control-0) was enriched in pathways for environmental sensing and stress response, including Two-component systems (4.78%) and ABC transporters (5.32%), reflecting an adaptive state for a fluctuating external environment (Fig. 7, Data S2). In contrast, spent Lees supplementation drove a consistent enrichment in core biosynthetic and metabolic pathways across all treatment groups by day 14. These included Ribosome biogenesis (11.41-13.24%), Aminoacyl-tRNA biosynthesis (2.26-2.46%), and the biosynthesis of essential amino acids such as Lysine (1.23-1.37%) and Histidine (1.29-1.52%). This biosynthetic shift was coupled with a dramatic reduction in pathways for the degradation of environmental pollutants and xenobiotics, such as Benzoate degradation, Aminobenzoate degradation, and Caprolactam degradation, which were prominent in the Control-0 group. The spent Lees-fed groups also showed a marked decrease in Bacterial chemotaxis and Flagellar assembly, indicating a lower investment in motility and environmental exploration. Furthermore, we observed an upregulation in pathways indicative of a sophisticated host-microbe interface.

**Fig. 7:**
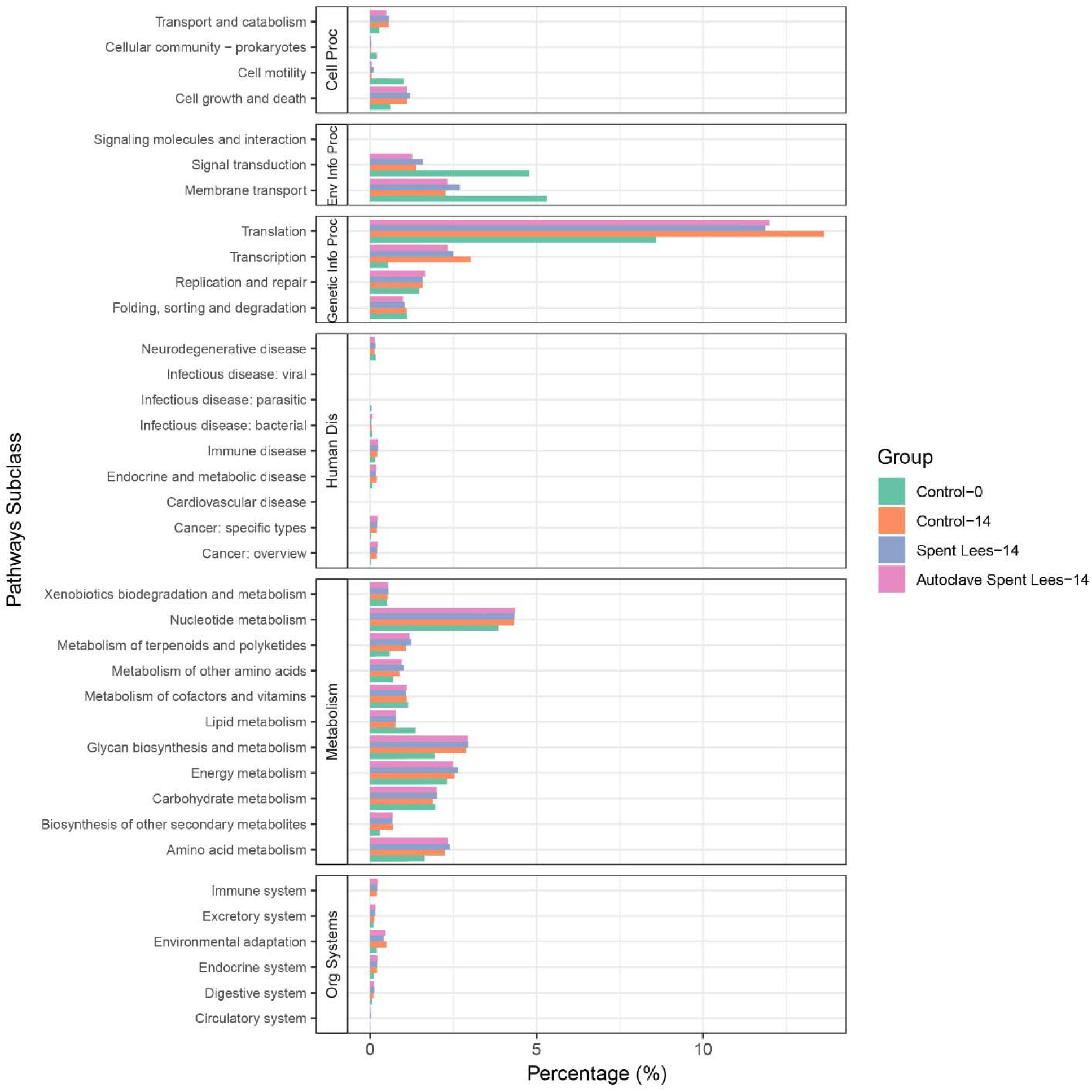
Functional metabolic landscape inferred from KEGG orthology. Stacked bar plots depict the relative contributions of major Kyoto Encyclopedia of Genes and Genomes (KEGG) pathway classes and subclasses across sample groups, derived from functional annotations of shotgun metagenomic datasets. Distinct colors denote individual functional categories, highlighting differences in metabolic capacity among groups. The principal pathway classes are abbreviated as follows: Cell Proc (Cellular Processes), Env Info Proc (Environmental Information Processing), Genetic Info Proc (Genetic Information Processing), Human Dis (Human Diseases), Metabolism, and Org Systems (Organismal Systems).

These included Lipopolysaccharide biosynthesis, essential for Gram-negative bacterial membrane integrity, and pathways for the metabolism of host-derived compounds, suggesting an enhanced capacity for interacting with and utilizing host resources. Dietary spent Lees orchestrates a fundamental metabolic transition in the gut microbiome. It suppresses the genetic machinery for environmental adaptation and pollutant detoxification, and instead promotes a community dedicated to robust protein synthesis, essential nutrient production, and stable host-microbe symbiosis. This functional reprogramming provides a mechanistic basis for the observed improvements in host growth and well-being.

## 4. Discussion

This study demonstrates that dietary spent Lees supplementation significantly enhances growth performance in poultry through a fundamental restructuring of the gut ecosystem. Our findings reveal that both spent Lees and autoclaved spent Lees function as powerful ecological levers, accelerating gut maturation and driving the microbiome toward a Bacteroides-dominated state that confers multiple benefits to the host.

### 4.1 Spent Lees-Driven Microbiome Maturation and the Bacteroides Dominance

The most striking effect of spent Lees supplementation was the dramatic ascendancy of *Bacteroides fragilis*, which reached 74.83% relative abundance in the spent Lees group. This represents a remarkable ecological shift from the early-life microbial community dominated by facultative anaerobes to a specialized, mature consortium. *Bacteroides* species possess expansive genetic arsenals for degrading complex polysaccharides^32^, making them particularly efficient at utilizing spent Lees components such as mannans and β-glucans as primary energy sources. The similar effect observed with autoclaved spent Lees confirms that viability is not essential for this prebiotic function, as the structural components remain biologically active after heat treatment^33^.

This spent Lees-driven enrichment of *B. fragilis* aligns with its established role as a keystone symbiont. Beyond its metabolic capabilities, *B. fragilis* modulates host immune responses through molecules like polysaccharide A^34^ and contributes to gut barrier integrity. Our network analysis positions *B. fragilis* as a central node within the “Mature Commensal Network,” strongly anti-correlated with the “Early-Life and Environmental Network” containing potential pathogens.

### 4.2 The Ecological Trade-off: Pathogen Exclusion Versus Diversity

The establishment of the Bacteroides-dominated community came with a significant ecological trade-off: markedly reduced microbial diversity, particularly in the spent Lees group. This finding challenges the conventional paradigm that higher diversity invariably indicates better health and necessitates a more nuanced interpretation of gut ecosystem states^35^. The benefits of this trade-off are substantial. The near-complete eradication of opportunistic pathogens like *Enterococcus faecium* and *Escherichia coli* represents a major improvement in gut health. These pathogens actively compete with the host for nutrients, damage intestinal epithelium, and trigger energy-intensive inflammatory responses^36^. Their exclusion likely redirects metabolic resources toward growth processes, directly contributing to the improved weight gain observed in spent Lees-supplemented birds. However, the diversity reduction involved suppression of beneficial short-chain fatty acid (SCFA) producers including *Faecalibacterium*, *Blautia*, and *Subdoligranulum*. Butyrate-producing bacteria like *Faecalibacterium prausnitzii* are crucial for colonic health, inflammation regulation, and serving as the primary energy source for colonocytes^37^. The graded suppression of these commensals—most severe with spent Lees—suggests that *B. fragilis’s* extreme fitness in the spent Lees-enriched environment may come at the expense of metabolic diversity. The superior final weight in the autoclaved spent Lees group, which maintained intermediate levels of SCFA producers, suggests that optimal production performance may require balancing pathogen exclusion with preservation of functional diversity. This balance may be context-dependent, with spent Lees potentially preferable in high-challenge environments and autoclaved spent Lees more suitable for optimal growth in cleaner conditions^38^.

### 4.3 Functional Benefits: Resistome and Virulome Restructuring

Beyond taxonomic shifts, spent Lees supplementation induced profound functional changes with significant implications for both animal and public health. The natural expansion of the resistome in control birds by day 14—characterized by high-risk genes including *tetW* and *o23S*—represents a concerning development in antimicrobial resistance (AMR) accumulation. Spent Lees supplementation substantially mitigated this expansion, particularly through the elimination of *o23S*. This resistome restructuring directly results from ecological changes. The correlation network analysis identified the “Early-Life and Environmental Network” as a major reservoir for broad-spectrum resistance genes^39^. By excluding this network, spent Lees supplementation removes a primary source of these genes. The mature, Bacteroides-dominated community that replaces it carries a different, less high-risk resistance profile^40^, resulting in net reduction of critical resistance determinants. Concurrently, spent Lees supplementation reshaped the gut virulome, significantly reducing genes encoding Type III Secretion System (T3SS) effectors—the “molecular syringes” used by pathogens like *E. coli* and *Salmonella* to inject effector proteins into host cells^41^. This suppression of offensive virulence mechanisms occurred alongside maintained or enriched genes for adherence and nutrient acquisition, indicating a shift in microbial strategy from host invasion toward commensal colonization^42^. This virulome reprogramming fosters a community optimized for peaceful persistence within the gut niche rather than aggressive host subversion^43^.

### 4.4 The Spent Lees Versus Autoclaved Spent Lees Paradox

The differential effects of spent Lees versus autoclaved spent Lees present a fascinating paradox. While spent Lees induced the most extreme microbial restructuring, autoclaved spent Lees yielded superior final weight. This divergence reflects distinct modes of action: spent Lees functions as both probiotic and prebiotic, while autoclaved spent Lees acts primarily as a prebiotic with enhanced nutritional delivery. Spent Lees actively interacts with the gut environment through pathogen sequestration, toxin degradation, and immune modulation^44^, creating a more competitive microenvironment that favors extreme *B. fragilis* dominance. In contrast, autoclaving lyses yeast cells in the spent Lees, releasing intracellular nutrients (nucleotides, peptides, vitamins) and increasing the accessibility of structural components^3^. This “passive” nutrient subsidy supports a slightly broader microbial consortium while directly providing the host with digestible protein, B-vitamins, and minerals^45^. The growth kinetics suggest spent Lees provides rapid early intervention against pathogens, while autoclaved spent Lees supports more sustained energy harvest through a balanced fermentative network. The metabolic specialization of *B. fragilis* toward acetate and succinate production, potentially at the expense of butyrate generation by other commensals^46^, might explain the slight long-term advantage of the more diverse autoclaved spent Lees community.

### 4.5 Integrated Mechanism: From Microbiome to Host Phenotype

Synthesizing our multi-omics data reveals a coherent mechanistic pathway from spent Lees supplementation to improved host phenotype: First, spent Lees introduction creates a new ecological niche based on complex polysaccharides, selectively enriching for specialized bacteria like *B. fragilis*^32^. Second, *B. fragilis* proliferation triggers ecological restructuring through competitive exclusion, suppressing early-life opportunists and their associated inflammatory potential^43^. Third, the established community shifts its functional profile toward host-beneficial biosynthesis—evidenced by enriched pathways for amino acid production and ribosome biogenesis^47^. Fourth, the combined effects of reduced pathogen competition, diminished inflammation, and enhanced microbial nutrient production free host metabolic resources for growth^48^.

### 4.6 Spent Lees as a Multi-Functional Supplement

Our findings support classifying spent Lees as a multi-functional supplement that simultaneously addresses gut health and nutrition. The prebiotic action of spent Lees components (MOS and β-glucans) selectively promotes beneficial commensals while blocking pathogen adhesion^49^, directly preventing dysbiosis. Concurrently, spent Lees provides high-quality protein, essential amino acids, B-vitamins, and minerals^50^, with autoclaving enhancing nutrient bioavailability^51^. These functions act synergistically: improved gut health enhances barrier function and nutrient absorption^52^, while spent Lees-derived B-vitamins support energy metabolism in both host and microbiota^53^. The microbial production of SCFAs from fermented spent Lees components provides additional energy for colonocytes, further supporting metabolic homeostasis^54^.

### 4.7 Study Limitations

While this study provides valuable insights into the effects of spent wash supplementation on poultry growth and gut microbiome ecology, several limitations warrant consideration. Most notably, we were unable to fully characterize the chemical composition of the spent lees, including the precise concentrations of specific bioactive compounds, nutrients, and potential contaminants, which limits our ability to establish direct structure-function relationships between specific components and the observed effects. The decision to pool fecal samples for metagenomic sequencing, while practical, precluded assessment of individual variation within treatment groups and limited statistical power for within-group comparisons. Furthermore, the 14-day experimental period, though sufficient to observe significant early changes, does not capture the long-term consequences of the dramatic ecological shifts observed, particularly regarding the sustainability of the monodominant *B. fragilis* state throughout a complete production cycle. Our functional analyses, while comprehensive in genetic potential assessment through metagenomics, lack validation through transcriptomic, proteomic, or metabolomic approaches to confirm whether these genetic changes translate to functional differences. The absence of pathogen challenge conditions, while demonstrating potent effects against natural microbiota, leaves the efficacy under deliberate disease exposure unverified. Additionally, the study focused primarily on microbial changes without complementary assessment of host physiological responses, including intestinal morphology, immune parameters, and barrier function. The use of a fixed supplementation level prevents determination of dose-response relationships, and the specific mechanisms underlying the remarkable selection for *B. fragilis* remain speculative. Finally, the assessment limited primarily to body weight metrics, without complementary measures of feed conversion ratio, carcass quality, or meat composition, provides an incomplete picture of production efficiency and product quality. These limitations, while not diminishing the significance of our findings, highlight important directions for future research to build upon this work.

## 5. Conclusion

This study demonstrates that dietary spent Lees supplementation significantly enhances poultry growth performance through fundamental restructuring of the gut ecosystem. Both spent Lees and autoclaved spent Lees formulations improved body weight gain, though through distinct mechanisms. Spent Lees drove more extreme microbiome changes, establishing remarkable *B. fragilis* monodominance and providing superior pathogen exclusion and antimicrobial resistance mitigation—completely eliminating the oxazolidinone resistance gene *o23S*. Autoclaved spent Lees yielded higher final weights, likely due to better preservation of beneficial SCFA-producers and enhanced nutrient delivery from lysed cells.

The benefits are substantial: reduced pathogenic loads, restructured resistome and virulome profiles, and enhanced metabolic capacity. However, significant trade-offs exist, particularly reduced microbial diversity and suppression of beneficial commensals like *Faecalibacterium* and *Lactobacillus*. The choice between spent Lees formulations should be context-dependent—spent Lees for pathogen-challenged environments where antimicrobial resistance mitigation is prioritized, and autoclaved spent Lees for optimal growth performance in controlled conditions.

Spent Lees supplementation represents a multi-functional approach to sustainable poultry production, simultaneously addressing growth performance, gut health, and antimicrobial resistance concerns. As a fermentation by-product, spent Lees offers an economically viable and environmentally sustainable alternative to conventional feed additives. Future research should optimize administration protocols and assess long-term ecosystem stability, but current evidence strongly supports spent Lees’s value as an effective alternative to antibiotic growth promoters in modern poultry production, while also contributing to waste valorization in fermentation industries.

## Acknowledgements

This work was supported by institutional resources from the Department of Microbiology and received partial support for reagents and consumables from the Research Cell, Jashore University of Science and Technology. The authors extend their sincere gratitude to Dr. Travis C. Glenn (Department of Environmental Health Science, University of Georgia, USA) for his valuable insights and critical review of the iNextEra library preparation protocol.

## Declaration of Generative AI in Writing

During the preparation of this work, the authors used DeepSeek (version 2.0) for language polishing and paraphrasing to improve readability. All content was thoroughly reviewed and edited by the authors, who take full responsibility for the publication’s content.

## Authors Contributions

MSR worked on spent lees collection, study design, monitoring, DNA extraction, sequencing data quality control, genomic and statistical analysis, presentation, and manuscript drafting. SI collected spent lees samples, conducted the farming study, collected the fecal samples, and assisted in drafting the manuscript. TJA assisted in farming study, DNA extraction, and manuscript drafting. AM and MRS assisted in DNA extraction, data analysis, presentation, and manuscript editing. MIUB assisted in study design, worked on Illumina library preparation and presentation, and critically reviewed the manuscript. SA monitored and led the overall study design, critically reviewed the manuscript, and finalized it.

## Data availability

The data that support the findings of this study are openly available in NCBI BioProject at https://www.ncbi.nlm.nih.gov/bioproject/ reference number PRJNA1355334.

## Code availability

All computational tools and scripts utilized in this study are available in a public GitHub repository: https://github.com/SShaminur/Microbiome-analysis-Pipeline.

---

**Supplementary Table S1:** Composition of Basal Feed collected from local market.

| Ingredients | Concentration (%) |
| --- | --- |
| Maize | 54.99 |
| Soya meal | 31 |
| Protein concentrate | 7 |
| Dicalcium phosphate | 1.35 |
| Limestone | 0.8 |
| Soybean oil | 4 |
| Lysine | 0.1 |
| Methionine | 0.12 |
| Vitamin premix | 0.25 |
| Choline chloride | 0.03 |
| Common salt | 0.36 |
| Total | 100 |

Supplementary Data S1: <u>Download link of Data S1</u>

Supplementary Data S2: <u>Download link of Data S2</u>

